# Validated B Vitamin Quantification from Lentils by Selected Reaction Monitoring Mass Spectrometry

**DOI:** 10.1101/2020.10.04.325548

**Authors:** Haixia Zhang, Devini De Silva, Dilanganie Dissanayaka, Thomas D. Warkentin, Albert Vandenberg

**Affiliations:** Crop Development Centre, Department of Plant Sciences, University of Saskatchewan, Saskatoon, S7N 5A8, Canada

**Keywords:** Lentil, quantification, B vitamins, germination, mass spectrometry, validation

## Abstract

A validated method for B vitamin separation and quantification from lentil seeds using ultra-performance liquid chromatography-selected reaction monitoring mass spectrometry (UPLC-SRM MS) was reported. The use of three enzymes (acid phosphatase, β-glucosidase, and rat serum) with a 4 h incubation was sufficient to convert bound B vitamins into their free forms. Twenty B vitamers were selected and a 5-min UPLC-SRM MS method was optimized for rapid analysis. This method was applied to quantify B vitamin concentration during lentil seed germination over a 5-day period. Total B vitamins increased up to 1.5-fold on day 5 (from 39.2 µg/g to 60.6 µg/g of dry weight) comparing with dry seeds. Vitamin B5 (pantothenic acid) was the most abundant B vitamin in both dry seeds (34.2%) and in germinated seeds (17.7%-24.5% of total B vitamins); B8 (biotin) and B12 (cyanocobalamin) were not detected in lentil samples.

## 1. Introduction

The B vitamin complex is a group of water-soluble vitamins which function as coenzymes in various biological processes in different organisms, including plants and humans. The B vitamin complex includes B1 (thiamine), B2 (riboflavin), B3 (niacin), B5 (pantothenic acid), B6 (pyridoxine), B8 (biotin), B9 (folic acid) and B12 (cyanocobalamin). Humans cannot synthesize B vitamins *in situ*, and any excess B vitamins in the body are eliminated through urine, therefore, humans require a continuous supply of B vitamins from their daily diet (Baj & Sieniawska, 2017).

Most B vitamins have multiple forms of vitamers which have similar chemical structure but different stability and unique function in metabolic pathways. B vitamin deficiency can lead to various chronic and acute diseases (Baj & Sieniawska, 2017) such as neural tube defects in the fetus (B9), anemia (B2) and beriberi (B1). Plant-based food can provide all of the essential B vitamins except for B12 (which is absent in higher plants). Pulses, such as lentil, chickpea, pea, common bean and faba bean, due to their high content of protein (>20%), micronutrients, dietary fiber and carbohydrate (Multari, Stewart, & Russell, 2015), have become one of the most promising staple foods. In Africa and the Middle East, pulses are the major food source of B vitamins.

B vitamins are present in free and chemically bound forms in plants or plant-based foods (Roje, 2007; Nurit, Lyan, Piquet, Branlard, & Pujos-Guillot, 2015). The heterogeneity of B vitamins greatly complicates their extraction and subsequent characterization. Existing B vitamin detection and quantification approaches (Zhang et al., 2018b; Fatima et al., 2019) include microbiological assays, gas chromatography (GC), capillary electrophoresis (CE), and liquid chromatography (LC), with UV, fluorescence, or mass spectrometry (MS) as detectors.

Procedures on B vitamin extraction from various foodstuffs were recently reviewed (Fatima et al, 2019; Zhang et al, 2018b). Most existing extraction methods were developed for pre-mixed vitamin supplements or processed foods, in which only a limited number of B vitamins in their free forms were extracted and characterized. For unprocessed foods, B vitamers are distributed within dense dry tissue and this requires fine grinding to make them accessible to the extraction solvents, and enzymes used to convert vitamers to their free B vitamin forms. Acid hydrolysis and subsequent enzyme treatment were commonly used (Fatima et al, 2019; Zhang et al., 2018b). Ndaw, Bergaentzlé, Aoudé-Werner & Hasselmann (2000) compared different extraction conditions of three B vitamers from various foods (pork, wheat flour and pea), and concluded that the use of acid hydrolysis is not necessary, and a mixture of enzymes (α-amylase, papain and acid phosphatase) was effective in releasing bound B vitamins in a single step. Nurit et al. (2015) studied the use of enzymes (acid phosphatase, taka-diastase and β-glucosidase) on B vitamin extraction from semi-coarse wheat flour products, and showed that the use of enzymes extracted higher amount of B vitamins, especially for nicotinamide (B3), riboflavin (B2) and thiamin (B1). Although promising, both reports required long enzyme hydrolysis duration (14-18 h), which is time consuming and low throughput. Leporati et al. (2005) reported three shorter extraction approaches for B vitamins from Italian pasta, with either 15 or 30 min extractions for B1, B2, B5 and B6, or 2 h incubation for folic acid (B9) extraction. Chatterjee et al. (2017) introduced an ultrasonication-assisted enzymatic extraction approach for analyzing nine B vitamers from fish, and reduced the enzyme hydrolysis to 4 h while maintaining good extraction efficiency.

In the last few years, the Vandenberg group developed 2-h enzyme hydrolysis approaches for folate (B9) extraction from seeds of pulses (Zhang, Jha, Warkentin, Vandenberg, & Purves, 2018a; Zhang et al., 2019), with quantification using UPLC-electrospray ionization mass spectrometry (ESI MS). The objective of the current work was to develop a simplified extraction workflow for B vitamin complex analysis from lentil seeds, and to establish and validate an efficient UPLC separation approach for subsequent quantification by ESI-MS.

## 2. Materials and methods

### 2.1. Chemical reagents and enzymes

2,3-dimercapto-1-propanol (BAL), 2-mercaptoethanol (MCE), 2-(N-morpholino)ehtanesulfonic acid (MES), activated charcoal, sodium ascorbate, protease (from *Streptomyces griseus*, PRO), α-amylase (from *Bacillus sp*., AMY), acid phosphatase (AP) and β-glucosidase (BGL) from almonds were purchased from Sigma-Aldrich (St. Louis, MO, USA). Dithiothreitol (DTT), potassium phosphate monobasic and dibasic, ammonium acetate and acetic acid were purchased from Fisher Scientific (Fair Lawn, NJ, USA). LC-MS grade acetonitrile and methanol (MeOH) was purchased form Thermo Fisher Scientific (Nepean, ON, Canada). Fresh rat serum (RS) was purchased from Lampire Biological Laboratories (Pipersville, PA, USA), shipped on wet ice, aliquoted into 10 mL upon arrival and stored at −80 °C before use. AP, BGL, AMY and PRO were prepared in water, with concentrations of 5 mg/mL for acid phosphatase and BGL, 4 mg/mL for PRO, and 20 mg/mL for AMY. Endogenous B vitamins in RS and PRO solution were removed by adding one-tenth (w/w) of activated charcoal as described previously (Zhang et al., 2018a), and were then stored at −80 °C before use.

### 2.2. Lentil seed germination

A zero-tannin lentil experimental line was selected for seed germination over a 5-day time course with 3 biological replicates (40 seeds per replicate). Seeds were rinsed with milliQ water to remove debris and dust particles, followed by rinsing with 70% ethanol for about 30 seconds. Seeds were then rinsed with milliQ water and placed on water-soaked filter papers in the plastic petri dishes for germination. The petri dishes were stored in a dark box at room temperature (21 °C) to start the germination process. A set of seeds after rinsing were immediately stored in −80 °C as the day 0 control. On day 1 (24 h after placing the seeds in the dishes), the germinating seeds were rinsed with milliQ water and transferred to falcon tubes for storage at −80 °C. This procedure was repeated for the subsequent four days. Three biological replicates were selected per day for up to 5 days. All the samples were freeze-dried on a lyophilizer (Labconco Corp., Kansas City, MO) for 30 h, ground into fine powder using a Udy grinder (UDY Corporation, Fort Collins, Colorado, USA) and stored at −80 °C before use.

### 2.3. B-vitamin standards and solution preparation

Isotope-labeled and non-labeled folate (B9) standards were obtained as reported previously (Zhang et al., 2018a; 2019). For the rest of the B vitamin standards, thiamine (B1), riboflavin (B2), nicotinic acid (B3), pantothenic acid (B5), biotin (B8), B6 (pyridoxine, pyridoxal and 4-pyridoxic acid), and cyanocobalamin (B12) were purchased from Sigma-Aldrich (St. Louis, MO, USA). Nicotinamide (B3) and pyridoxamine (B6) were from Fisher Scientific (Waltham, MA, USA). ^13^C_4_ ^15^N_2_−riboflavin, ^13^C_3_ ^15^N-nicotinamide, ^13^C_3_ ^15^N-pantothenic acid, D_4_ -biotin, D_3_ - pyridoxine and ^13^C_3_-thiamine were from Cambridge Isotope Laboratories (Tewksbury, MA).

Folates (B9) standard solutions were prepared as described previously (Zhang et al., 2019). For the rest of the B vitamins, stock solutions of 1.0 mg/mL were prepared in 50% acetonitrile (Nurit et al., 2015), and the working solutions (0.1 or 0.5 mg/mL) were diluted from the stock solution with water. All solutions were aliquoted and stored at −80 °C before use.

### 2.4. Extraction of B vitamins

B-vitamin extraction was based on previous work (Zhang et al., 2018a, 2019) on folate extraction from seeds of pulses, and two other reports (Ndaw et al., 2000; Nurit et al., 2015) with some modifications. Degassed Milli-Q water (Millipore, Milford, MA, USA) was used to prepare all buffer solutions. The extraction buffer was 50 mM MES (pH 6). Different antioxidants were added to the extraction buffer to examine their effect on B vitamin stability, and were specified in the later section. For method optimization, a common lentil cultivar, CDC Maxim, was used as a model cultivar with three technical replicates used for each extraction condition. In general, 30 mg (± 1.0 mg) finely ground seed powder was weighed into a 2 mL microvial (Sarstedt, Germany). To each vial, 450 μL extraction buffer containing nine stable isotope labeled internal standards was added (^13^C_5_-folic acid, ^13^C_5_-5-methyltetrahydrofolate (^13^C_5_-5-MTHF), ^13^C_5_-5-formyltetrahydrofolate (^13^C_5_-5-FTHF), ^13^C_4_^15^N_2_-riboflavin, ^13^C_3_15N-nicotinamide, ^13^C_3_ ^15^N-pantothenic acid, D_4_ -biotin, D_3_ -pyridoxine and ^13^C_3_ -thiamine). Samples were vortexed and heated for 5 min at 100 °C with shaking (1200 rpm) on the Vortemp 56 shaking incubator (Montreal-Biotech Inc., Kirkland, QC). Samples were cooled on ice, and different combination of enzymes were added. These enzymes were RS, AP, BGL, PRO and AMY, and the final volume was adjusted to 900 µL by addition of extraction buffer (without internal standards). Different incubation times was also examined for enzyme hydrolysis. The final optimized one-step extraction procedure is summarized here. 50 mM MES (pH 6.0) solution containing 1% (w/v) sodium ascorbate and 0.2% (v/v) MCE was used as the extraction buffer. For 30 mg of seed powder, 450 µL of extraction buffer containing nine internal standards (Table 2), and 170 μL extraction buffer alone were added and vortexed for 6 s. The mixture was heated for 5 min at 100 °C, after cooling on ice, 120 μL RS, 80 μL AP, and 80 μL BGL solution were added and vortexed for 10 s. Samples were incubated for 4 h at 37 °C with shaking at 1200 rpm. Samples were cooled on ice and centrifuged at 4 °C and 16,600 g for 30 min. 200 μL supernatant was transferred to 10 kDa MWCO filters (PALL, New York, NY, USA), and centrifuged as above. The filtrates were directly transferred to glass inserts inside amber vials for LC-SRM analysis. All the experiments were conducted in a fume hood under dim light. Three technical replicates were conducted per condition, and two method blank vials with no samples were also prepared for each analysis. Each extract was injected twice for data collection. For germinated lentils, three biological replicates were selected, and each extract was also injected twice on LC-MS.

### 2.5. Ultra-performance liquid chromatography selected reaction monitoring mass spectrometry (UPLC-SRM MS) analysis

B-vitamin separation and detection was conducted on a Thermo Fisher Vanquish− Flex UPLC combined with TSQ Altis electrospray ionization (ESI) triple quadrupole mass spectrometer operated in positive ion mode (San Jose, CA, USA). Twenty non-labeled and 9 isotope labeled B vitamers were included in this study, and the fragmentation of individual B vitamer standard was optimized by direct infusion, in a solvent of 50% acetonitrile containing 0.1% acetic acid. One fragment ion per B vitamer was selected for subsequent LC-SRM analysis, and the corresponding collision energy and RF-lens values of these B vitamers are listed in **Table 1**.

**Table 1.** The UPLC-SRM MS parameters of B vitamers and internal standards.

| Compounds | Chemical formula | Molecular mass | Retention Time (min) | Precursor (m/z) | Product (m/z) | Collision Energy (V) | RF Lens (V) | Quantification? |
| --- | --- | --- | --- | --- | --- | --- | --- | --- |
| Thiamine (B1) | C<sub>12</sub>H<sub>17</sub>N<sub>4</sub>OS | 265.4 | 2.55 | 265.1 | 122.1 | 14.7 | 30 | Y |
| Riboflavin (B2) | C<sub>17</sub>H<sub>20</sub>N<sub>4</sub>O<sub>6</sub> | 376.4 | 3.38 | 377.2 | 172.1 | 37.0 | 91 | Y |
| Nicotinamide (B3) | C<sub>6</sub>H<sub>6</sub>N<sub>2</sub>O | 122.1 | 2.93 | 123.1 | 78.1 | 23.1 | 56 | Y |
| Nicotinic acid (B3) | C<sub>6</sub>H<sub>5</sub>NO<sub>2</sub> | 123.1 | 1.94 | 124.1 | 78.1 | 22.3 | 64 | Y |
| Pantothenic acid (B5) | C<sub>9</sub>H<sub>17</sub>NO<sub>5</sub> | 219.2 | 2.16 | 220.1 | 90.1 | 13.5 | 30 | Y |
| Pyridoxamine (B6) | C<sub>8</sub>H<sub>12</sub>N<sub>2</sub>O<sub>2</sub> | 168.2 | 1.31 | 169.1 | 77.1 | 36.0 | 32 | Y |
| Pyridoxine (B6) | C<sub>8</sub>H<sub>11</sub>NO<sub>3</sub> | 169.2 | 2.25 | 170.1 | 77.1 | 35.3 | 46 | Y |
| Pyridoxal (B6) | C<sub>8</sub>H<sub>9</sub>NO<sub>3</sub> | 167.2 | 2.27 | 168.1 | 94.1 | 23.3 | 30 | Y |
| 4-pyridoxic acid (B6) | C<sub>8</sub>H<sub>9</sub>NO<sub>4</sub> | 183.2 | 2.88 | 184.1 | 65.1 | 33.5 | 36 | Y |
| Biotin (B8) | C<sub>10</sub>H<sub>16</sub>N<sub>2</sub>O<sub>3</sub>S | 244.3 | 3.18 | 245.1 | 97.1 | 31.2 | 42 | Y |
| 5,10-methenyl THF (B9) | C<sub>20</sub>H<sub>21</sub>N<sub>7</sub>O<sub>6</sub> | 455.4 | 2.77 | 456.2 | 412.3 | 29.3 | 190 | Y |
| THF (B9) | C<sub>19</sub>H<sub>23</sub>N<sub>7</sub>O<sub>6</sub> | 445.4 | 2.94 | 446.2 | 299.3 | 19.4 | 63 | Y |
| 10-FFA (B9) | C<sub>20</sub>H<sub>19</sub>N<sub>7</sub>O<sub>7</sub> | 469.4 | 3.07 | 470.2 | 295.2 | 24.6 | 80 | Y |
| 5-FTHF (B9) | C<sub>20</sub>H<sub>23</sub>N<sub>7</sub>O<sub>7</sub> | 473.4 | 3.11 | 474.2 | 299.3 | 31.3 | 62 | Y |
| Folic acid (B9) | C<sub>19</sub>H<sub>19</sub>N<sub>7</sub>O<sub>6</sub> | 441.4 | 3.17 | 442.2 | 295.2 | 16.0 | 53 | Y |
| 5-MTHF (B9) | C<sub>20</sub>H<sub>25</sub>N<sub>7</sub>O<sub>6</sub> | 459.5 | 3.22 | 460.2 | 313.3 | 19.0 | 65 | Y |
| 7,8-DHF (B9) | C<sub>19</sub>H<sub>21</sub>N<sub>7</sub>O<sub>6</sub> | 443.4 | 3.23 | 444.2 | 178.2 | 10.2 | 52 | Y |
| 10-FDHF (B9) | C<sub>20</sub>H<sub>21</sub>N<sub>7</sub>O<sub>7</sub> | 471.4 | 2.90 | 472.2 | 178.2 | 22.2 | 123 | Y |
| Cyanocobalamin (B12) | C<sub>63</sub>H<sub>88</sub>CoN<sub>14</sub>O<sub>14</sub>P | 1355.4 | 3.23 | 678.4 | 147.2 | 42.7 | 113 | Y |
| MeFox | C<sub>20</sub>H<sub>23</sub>N<sub>7</sub>O<sub>7</sub> | 473.4 | 2.63 | 474.2 | 284.2 | 37.0 | 78 | Y |
| <sup>13</sup>C<sub>5</sub>-5-MTHF | <sup>13</sup>C<sub>5</sub>C<sub>15</sub>H<sub>25</sub>N<sub>7</sub>O<sub>6</sub> | 464.5 | 3.22 | 465.2 | 313.3 | 18.9 | 62 | IS |
| <sup>13</sup>C<sub>5</sub>-folic acid | <sup>13</sup>C<sub>5</sub>C<sub>14</sub>H<sub>19</sub>N<sub>7</sub>O<sub>6</sub> | 445.4 | 3.18 | 447.2 | 295.2 | 15.6 | 52 | IS |
| <sup>13</sup>C<sub>5</sub>-5-FTHF | <sup>13</sup>C<sub>5</sub>C<sub>15</sub>H<sub>23</sub>N<sub>7</sub>O<sub>7</sub> | 478.4 | 3.11 | 479.2 | 327.3 | 19.0 | 62 | IS |
| <sup>13</sup>C<sub>3</sub>-thiamine | <sup>13</sup>C<sub>3</sub>C<sub>9</sub>H<sub>17</sub>N<sub>4</sub>OS | 268.4 | 2.56 | 268.1 | 122.1 | 14.5 | 30 | IS |
| <sup>13</sup>C<sub>3</sub><sup>15</sup>N-pantothenic acid | <sup>13</sup>C<sub>3</sub>C<sub>6</sub>H<sub>17</sub><sup>15</sup>NO<sub>5</sub> | 223.2 | 2.17 | 224.1 | 94.1 | 14.0 | 35 | IS |
| D<sub>3</sub>-pyridoxine | C<sub>8</sub>D<sub>3</sub>H<sub>8</sub>NO<sub>3</sub> | 172.2 | 2.21 | 173.1 | 136.2 | 21.2 | 37 | IS |
| D<sub>4</sub>-biotin | C<sub>10</sub>D<sub>4</sub>H<sub>12</sub>N<sub>2</sub>O<sub>3</sub>S | 248.3 | 3.17 | 249.1 | 170.2 | 22.2 | 37 | IS |
| <sup>13</sup>C<sub>3</sub><sup>15</sup>N-nicotinamide | <sup>13</sup>C<sub>3</sub>C<sub>3</sub>H<sub>6</sub><sup>15</sup>NNO | 126.1 | 2.92 | 127.1 | 83.1 | 21.0 | 61 | IS |
| <sup>13</sup>C<sub>4</sub><sup>15</sup>N<sub>2</sub>-riboflavin | <sup>13</sup>C<sub>4</sub>C<sub>13</sub>H<sub>20</sub><sup>15</sup>N<sub>2</sub>N<sub>2</sub>O<sub>6</sub> | 382.4 | 3.38 | 383.1 | 249.1 | 22.0 | 93 | IS |

5,10-methenyl THF, 5,10-methenyltetrahydrofolate; THF, 5,6,7,8-tetrahydrofolate; 10-FFA, 10-formyl folic acid; 5-FTHF, 5-formyltetrahydrofolate; 5-MTHF, 5-methyltetrahydrofolate; 7,8-DHF, 7,8-dihydrofolate; 10-FDHF, 10-formyldihydrofolate; MeFox, pyrazino-*s*-triazine derivative of 4α-hydroxy-5-methyltetrahydrofolate. IS, internal standard.

The performance of several LC columns and different LC solvents were examined and will be specified in the Discussion section. The final LC condition was as follows: an Agilent ZORBAX SB-Aq column (100 mm × 2.1 mm, 1.8 μm particle size) with a SB-Aq guard column (2.1 × 5 mm, 1.8 um particle size) was selected, and the mobile phase consisted of 10 mM ammonium acetate and 0.1% acetic acid in water (solvent A) or in MeOH (solvent B). The flow rate was set to 0.3 mL/min and a 5-min gradient was used. The gradient started with 5% solvent B and maintained for 0.5 min, solvent B was ramped to 15% at 0.6 min and increased to 90% at 3.0 min, it then was maintained for an additional 1 min, and returned to 5% at 4.1 min, and maintained for 0.9 min to equilibrate the column. The LC column temperature was controlled at 25 °C and the autosampler at 6 °C. To avoid direct light, the light in the autosampler was turned off and the autosampler door was shielded. The mass spectrometer parameters were: electrospray voltage 3000 V, vaporizer temperature 350 °C, sheath gas pressure 60, auxiliary gas pressure 10, sweep gas 1, and capillary temperature 325 °C.

### 2.7. Method validation

Method validation was based on previous reports (Zhang et al., 2019) with some modifications. Parameters to validate include sensitivity (limit of detection, LOD; and limit of quantification, LOQ), linearity (R^2^, linear range), accuracy, precision, recovery and matrix effect. A blank matrix solution was prepared from CDC Maxim. In brief, 30 ml of 50 mM MES buffer (pH 6.0) was added to 1 g of seed powder, the mixture was heated 10 min in a boiling water bath. The solution was cooled to room temperature, and 3.3 g activated charcoal was added, for 1 h incubation at room temperature, with occasionally mixing in between. The mixture was then aliquoted into 2 mL microtubes and centrifuged for 30 min at 20,000 g, the supernatant was pooled and filtered through a 0.2 um syringe filter. The presence of endogenous B vitamins was tested using established LC-SRM MS method. Thiamine and pantothenic acid cannot be removed effectively, and were present in low amount in the treated matrix. Their signals were deducted from the method blank before further analysis. The treated matrix was stored at −80 °C before use.

Three sets of four calibration curves were prepared according to previous reports (Zhang, 2018a; 2019), including the method blank which containing the internal standards. In brief, set A was prepared in MES buffer, set B was prepared in charcoal treated matrix, and both sets of curves were injected for UPLC-SRM analysis after preparation. Set C was prepared in treated matrix and went through the entire extraction procedure, that is, heated for 5 min at 100 °C, cooled to room temperature, and then three enzymes were added (rat serum, BGL, and AP) and incubated for 4 h at 37 °C with mixing (1100 rpm). The solution was centrifuged for 30 min at 20,000 g, 200 µl of the supernatant was transferred out for filtering through 10 kD MWCO, the final filtrate was injected for UPLC-SRM analysis. For each set, eight-point calibration curves were prepared with series dilution, and nine isotope labeled internal standards (Table 2) were added at constant concentration. 1% sodium ascorbate and 0.2% MCE were added to each solution before use. Three quality control (QC) solutions were also prepared for accuracy and precision measurements, and their concentrations were listed in Table 3. LOD, LOQ, recovery, and matrix effect was determined as reported previously (Zhang et al., 2018a; 2019).

**Table 2.**
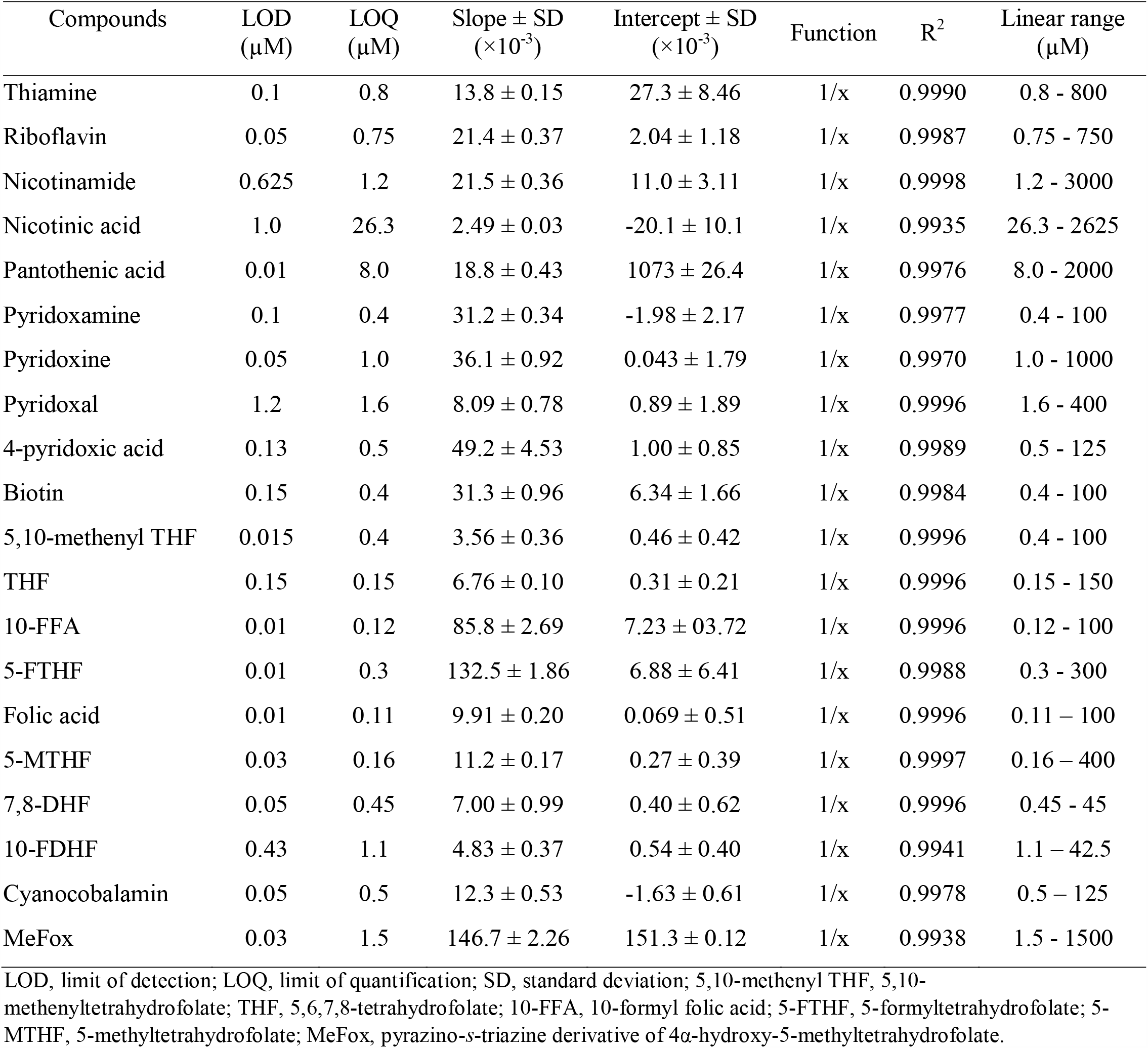
Calibration curve and sensitivity data for B vitamers and MeFox (n=4); calibration curves were prepared in treated lentil matrix (set B).

**Table 3.** Measurement accuracy (%) of three QC standards (low, medium, and high concentrations) from three set of calibration curves. Data were from the average of six replicates (3 biological × 2 analytical replicates).

| Compounds | Set A |  |  | Set B |  |  | Set C |  |  | QC concentrations,<br>low to high (nM) |
| --- | --- | --- | --- | --- | --- | --- | --- | --- | --- | --- |
|  | low | medium | high | low | medium | high | low | medium | high |  |
| Thiamin | 84.0 | 92.4 | 93.6 | 83.1 | 95.3 | 96.8 | NA | 94.4 | 95.9 | 4.7, 35.6, 266.7 |
| Riboflavin | 86.5 | 86.2 | 94.6 | 89.4 | 85.3 | 93.6 | 100.4 | 90.6 | 95.5 | 4.4, 33.3, 250 |
| Nicotinamide | 92.2 | 93.3 | 94.7 | 88.5 | 94.9 | 96.8 | 91.2 | 94.6 | 96.6 | 17.8, 133.3, 1000 |
| Nicotinic acid | NA | 68.4 | 77.4 | NA | 74.5 | 83.8 | NA | 82.7 | 87.1 | 15.6, 116.7, 875 |
| Pantothenic acid | 77.9 | 85.0 | 87.5 | 85.8 | 90.0 | 90.8 | 83.9 | 90.2 | 92.3 | 11.9, 88.9, 666.7 |
| Pyridoxamine | 81.7 | 78.8 | 86.3 | 74.7 | 81.3 | 88.8 | 121.7 | 89.1 | 86.6 | 0.6, 4.4, 33.3 |
| Pyridoxine | 85.8 | 87.5 | 93.6 | 88.8 | 89.7 | 93.8 | 87.9 | 91.0 | 93.1 | 5.9, 44.4, 333.3 |
| Pyridoxal | 85.8 | 92.7 | 94.7 | 79.4 | 90.3 | 95.4 | NA | 92.9 | 92.6 | 2.4, 17.8, 133.3 |
| 4-pyridoxic acid | 83.6 | 79.9 | 89.9 | 96.1 | 85.7 | 91.8 | NA | 101.2 | 91.8 | 0.7, 5.6, 41.7 |
| Biotin | 99.5 | 97.6 | 89.9 | 72.5 | 94.1 | 95.3 | NA | 99.5 | 97.9 | 0.6, 4.4, 33.3 |
| 5,10-methenyl THF | 88.6 | 93.2 | 88.5 | 93.7 | 95.4 | 96.6 | 81.8 | 79.6 | 85.9 | 0.6, 4.4, 33.3 |
| THF | 90.6 | 95.0 | 93.0 | 93.8 | 92.3 | 95.8 | 92.7 | 93.8 | 93.4 | 0.9, 6.7, 50 |
| 10-FFA | 103.9 | 93.8 | 92.1 | 106.1 | 92.1 | 95.7 | 104.2 | 94.5 | 92.5 | 0.7, 5.3, 39.6 |
| 5-FTHF | 89.0 | 88.9 | 92.5 | 96.1 | 86.8 | 94.6 | 89.6 | 92.9 | 92.4 | 1.8, 13.3, 100 |
| Folic acid | 87.2 | 94.9 | 94.6 | 104.6 | 93.9 | 93.2 | 80.6 | 98.0 | 92.7 | 0.7, 5.0, 37.5 |
| 5-MTHF | 89.5 | 89.0 | 93.9 | 88.4 | 89.2 | 94.9 | 94.8 | 90.7 | 93.4 | 2.4, 17.8, 133.3 |
| 7,8-DHF | 95.3 | 97.5 | 95.4 | 86.0 | 94.4 | 93.7 | 77.6 | 102.5 | 98.2 | 0.7, 5.0, 37.5 |
| 10-FDHF | NA | 108.3 | 102.8 | NA | 103.5 | 94.8 | NA | 84.1 | 91.6 | 0.6, 4.7, 35.4 |
| Cyanocobalamin | 92.9 | 93.4 | 95.1 | 83.1 | 91.2 | 91.6 | NA | 109.0 | 98.9 | 0.7, 5.6, 41.7 |
| MeFox | 93.8 | 107.8 | 99.7 | 104.2 | 109.8 | 105.8 | 86.8 | 94.6 | 99.3 | 8.9, 66.7, 500 |

Accuracy (%) is calculated as average observed concentration/nominal concentration × 100%. NA, not available, as these concentrations were below the limit of quantification. 5,10-methenyl THF, 5,10-methenyltetrahydrofolate; THF, 5,6,7,8-tetrahydrofolate; 10-FFA, 10-formyl folic acid; 5-FTHF, 5-formyltetrahydrofolate; 5-MTHF, 5-methyltetrahydrofolate; 7,8-DHF, 7,8-dihydrofolate; 10-FDHF, 10-formyldihydrofolate; MeFox, pyrazino-*s*-triazine derivative of 4α-hydroxy-5-methyltetrahydrofolate.

### 2.7. Calibration curve and concentration determination

For the final calibration curve set up, nine isotope labelled internal standards were used (listed in Table 2), and calibration curves were built from 20 B vitamin standards, thiamine, riboflavin, nicotinamide, nicotinic acid, pantothenic acid, biotin, pyridoxine, pyridoxal, pyridoxamine, 4-pyridoxic acid, cyanocobalamin, folic acid, 5,6,7,8-tetrahydrofolate (THF), 5-MTHF, 5-FTHF, 5,10-methenyltetrahydrofolate (5,10-methenyl THF), 10-formylfolic acid (10-FFA), 10-formyldihydrofolate (10-FDHF), 7,8-dihydrofolate (7,8-DHF), and a pyrazino-*s*-triazine derivative of 4α-hydroxy-5-methyltetrahydrofolate (MeFox). Besides working as internal standards for their own corresponding B vitamers, the isotope labelled internal standards were also used to normalize other vitamers which had no isotope labeled counterparts. For example, ^13^C_5_-folic acid was used to normalize THF, 7,8-DHF and 10-FFA; ^13^C_5_-5-MTHF for 5,10-methenyl THF and MeFox; ^13^C_5_-5-FTHF for 10-FDHF, ^13^C_3_ ^15^N-nicotinamide for nicotinic acid, D_3_-pyridoxine for pyridoxal, pyridoxamine and 4-pyridoxic acid; and ^13^C_4_ ^15^N_2_-riboflavin was used for cyanocobalamin.

The concentration range of each standard was determined in a batch of preliminary experiments. An eight-point calibration curve was built, and a method blank including internal standards and enzymes but not lentil samples was used for comparison, and a 1/x weighted linear regression was used for all vitamers. Due to the instability of some B vitamers, calibration curve solutions were prepared on the same day as the samples, and were injected at the beginning, in the middle, and at the end of the queue. The peak area ratios of the target B vitamers to their corresponding internal standards were used to build the calibration curves.

## 3. Results and discussions

### 3.1. Optimization of B vitamin separation on UPLC

B vitamin separation and quantification using liquid chromatography have been frequently reported, with various number of B vitamers being simultaneously analyzed (Roelofsen-de Beer, van Zelst, Wardleb, Kooij, & de Rijke, 2017; Asante et al., 2018). To date, the most comprehensive method quantified 13 B vitamers using a 40 min LC method with a diode matrix detector for peak detection (Bendryshev, Pashkova, Pirogov, & Shpigun, 2010). To take advantage of the high selectivity of a triple quadruple mass spectrometer, we decided to develop a fast UPLC-SRM MS approach for B vitamin analysis.

Twenty B vitamers from eight B vitamin families were selected for LC method development. They are B1 (thiamine), B2 (riboflavin), B3 (nicotinamide, nicotinic acid), B5 (pantothenic acid), B6 (pyridoxine, pyridoxamine, pyridoxal and 4-pyridoxic acid), B8 (biotin), B9 (THF, 5-MTHF, 5,10-methenyl THF, 10-FFA, 5-FTHF, FA, 7,8-DHF, 10-FDHF, and a microbiologically inactive folate derivative, MeFox), and B12 (cyanocobalamin). A collection of UPLC columns were evaluated for their separation efficiency, these included Waters HSS T3 (150 × 2.1 mm, 1.8 µm particle size) (Jha et al., 2015; Zhang et al., 2018a, 2019), Waters BEH C18 (2.1 × 100 mm, 1.7 µm), Agilent Eclipse pulse C18 (2.1 × 50 mm, 1.8 µm), Agilent ZORBAX SB-Aq with two different lengths (ID 2.1 mm, 50 or 100 mm in length, 1.8 µm), and Agilent ZORBAX Extend C18 (2.1 × 100 mm, 1.8 µm). Organic solvent such as acetonitrile and methanol were tested as LC elution solvent (solvent B), and different concentrations of ammonium acetate (0, 5, 10 and 20 mM) in combination of 0.1% acetic acid were studied as solvent additive. Different lengths of LC gradients were also investigated, at 15 min, 10 min, and 5 min separately. Overall, the Agilent ZORBAX SB-Aq column with the length of 100 mm had the best performance compared with other LC columns (data not shown), methanol was selected as the elution solvent (solvent B), 10 mM ammonium acetate and 0.1% acetic acid in both solvents resulted in the best peak shape, and a 5-min LC gradient could provide good separation of the target analytes while maintaining short analysis time. The final optimized LC method and the solvent gradient was described in the material and method section, and the retention time of each B vitamer was listed in Table 2.

### 3.2. Method development on B vitamin extraction

One problem which hinders B vitamin extraction and subsequent analysis is their instability under various conditions (Fatima et al, 2019), such as UV light (photodegradation), air (oxidation), temperature and storage conditions (solution pH, storage time). The stability and interconversion of vitamin B9 (folates) are well studied and documented (De Brouwer, Zhang, Storozhenko, Van Der Straeten, & Lambert, 2007; Zhang et al., 2018a; Strandler, Patring, Jägerstad, & Jastrebova, 2015), therefore antioxidants are always used for folate analysis (Patring et al., 2005). For the other B vitamins, antioxidant reagents were rarely used in previous reports. Since the aim of the current work was to analyze as many vitamers as possible, we tested the effect of antioxidants on B vitamin extraction in a preliminary study (data not shown), using CDC Maxim as the model lentil seeds, and 1% sodium ascorbate together with either 0.5% DTT, 0.2% MCE or 0.2% BAL as the antioxidation reagents (Zhang et al., 2019). It was observed that the addition of antioxidants could improve the stability of some of the B vitamers, such as 4-pyridoxic acid, pyridoxal, THF, and 5,10-methenyl THF. When combined with sodium ascorbate, DTT, MCE, and BAL had similar antioxidation effects. Since MCE is in liquid form and easy to handle, 1% sodium ascorbate and 0.2% MCE were used in the extraction buffer for further method optimization.

Enzyme selection to release bound B vitamins was based on previous reports. Ndaw et al. (2000) conducted an analysis of B1, B2 and B6 from various foodstuffs such as rice and pea, and suggested that the use of an enzyme mixture (α-amylase, papain and acid phosphatase) in a single step hydrolysis was an efficient way to release bound B vitamins. Nurit et al. (2015) employed a four-enzyme mixture (acid phosphatase, papain, taka-diastase and BGL) for B vitamin (B1, B2, B3, B5 and B6) extraction out of wheat flour products. Jha et al. (2015) initiated folate (B9) extraction using a tri-enzyme, three-step hydrolysis approach, then they proposed a two-enzyme, single-step folate extraction approach (Zhang et al., 2018a), and later reported the use of a single-source of enzyme (rat serum) was as effective as previous work (Zhang et al., 2019).

In the current work, for complete extraction of all B vitamins out of lentil seeds, three essential enzymes were selected, RS, AP and BGL. RS functions to convert polyglutamate folates (B9) into monoglutamate folates (Zhang et al., 2019), AP is for dephosphorylation and converting B1, B2, B6 into their free forms (Ndaw et al., 2000), and BGL is required to determine the glycosylated forms of vitamin B6 (van den Berg et al., 1996). The use of AMY and PRO on B vitamin extraction from lentils was examined herein, in combination of these three essential enzymes. Considering the optimum pH of each enzyme, AP (pH 4-7), BGL (pH 5), PRO (pH 5-9), AMY (pH 6.9), and RS (pH 6-8), a 50 mM MES solution (pH 6.0) was used as the extraction buffer.

In addition, the level of endogenous B vitamins from these enzymes were also examined. Acid phosphatase, BGL or α-amylase solution had no detectable B vitamins, while protease and rat serum, after charcoal treatment (Zhang et al., 2019), still had detectable pyridoxal, riboflavin, pantothenic acid, thiamine and biotin respectively. Therefore for later sample analysis, a method blank was used to measure the endogenous levels of aforementioned B vitamers in the extraction solution, and values determined were deducted from the sample data.

The total B vitamins extracted from CDC Maxim using different combination of enzymes and under various incubation periods (1 h, 2 h, 4 h, and overnight, respectively, at 37 °C) are shown in Figure 1. Cyanocobalamin (B12) and biotin (B8) were below the limit of quantification in CDC Maxim, and thus were not listed in this figure. Using results from the three-enzyme incubation as control, at each time point, the addition of α-amylase, protease or both of them resulted in similar amount of total B vitamins at each time point, with 4 h incubation giving the highest total B vitamins (42.6 µg/g for using three enzymes, 41.4 µg/g with addition of a-amylase, 42.3 µg/g with addition of protease, and 41.5 µg/g using five enzymes). Overnight incubation led to the lowest total amount (8.6% to 20.4% decrease of total B vitamins comparing to 4 h incubation). This is mainly due to the gradual release of B3 (nicotinic acid and nicotinamide) within the first four hours incubation (increased 17% to 54% comparing to those from 1 h incubation), and its significant loss during overnight incubation (reduced 69% to 98% comparing to those from 4 h incubation, (Figure 1). A similar observation was reported by Chatterjee et al. (2017), with a 4 h extraction approach resulting in a higher level of B3 (nicotinic acid and nicotinamide) in fish comparing to 18 h incubation. A recent report on B vitamin quantification from faba beans seeds (Marshall, Zhang, Kateil, & Vandenberg, 2020) indicated that an incubation time between 2 to 4 h was a good choice, whereas for the current lentil sample analysis, 4 h incubation gave slightly higher concentration of total B vitamins (5.8%-10.3% higher) than those from 2 h incubation. This could be due to the higher proportion of seed coat in lentil comparing to that in faba bean, since seed coat is mainly composed of cellulose which is slow to hydrolyze, and thus leads to slow release of B vitamins. Note that since biotin (B8) or cyanocobalamin (B12) were not detected in lentil seeds, for samples containing these two B vitamins, the extraction conditions may need to be optimized accordingly.

**Figure 1.**
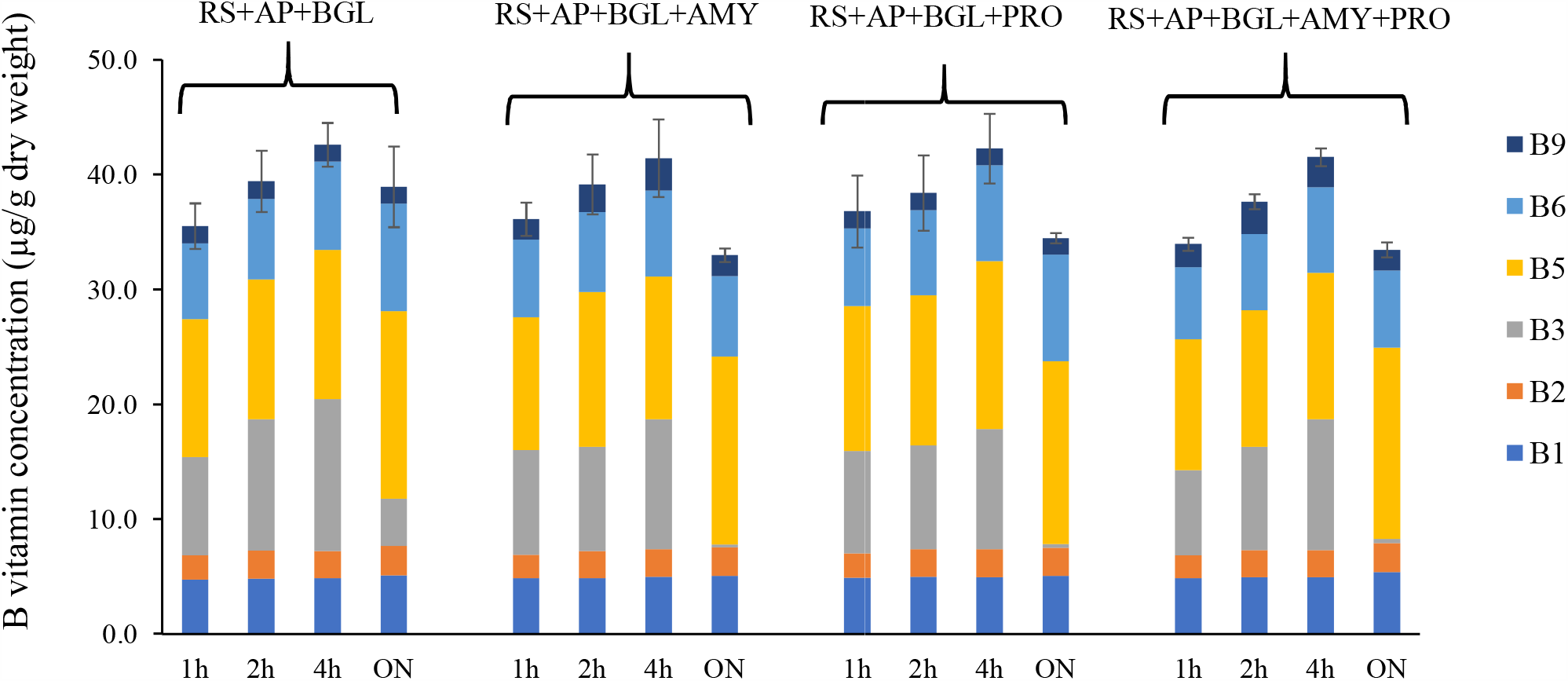
Total B vitamins extracted from CDC Maxim (lentil), with different combination of enzymes and under various incubation period. Extraction buffer was pH 6.0, 50 mM MES containing 1% sodium ascorbate and 0.2% MCE, incubation temperature was 37 °C. RS, rat serum; AP, acid phosphatase; BGL, β-glucosidase; AMY, α-amylase; PRO, protease; ON, overnight extraction. Error bars were the standard deviation from six replicates (3 technical × 2 analytical replicates).

Since the addition of α-amylase or protease did not show significant improvement of B vitamin extraction, in order to simplify the extraction procedure, the use of three enzymes (rat serum, acid phosphatase, and β-glucosidase) and a 4 h incubation at 37 °C was employed for further study.

### 3.3. Method validation

The final method was validated based on previous reports (Zhang et al., 2019), with a blank matrix prepared from a common lentil cultivar, CDC Maxim, as described in the Materials and Methods section. Three sets of calibration curves were prepared either in extraction buffer (set A), or in treated lentil matrix (set B and C). Curves from set A and B were injected immediately for LC-MS after preparation, whereas set C curves went through the entire extraction procedure (heated for 5 min at 100 °C, three enzymes were added and then incubated for 4 h at 37 °C, centrifuged and filtrated through MWCO filters) before LC injection. Recognizing that some of the endogenous B vitamins, such as thiamine, pantothenic acid, riboflavin, biotin and pyridoxal cannot be completely removed from treated matrix or enzyme solution, method blanks were prepared for all calibration curves, and signals from matrix blanks were deducted before further validation. The calibration curve parameters and sensitivity data for each B vitamer from set B were listed in Table 2. The measurement accuracy of three QC samples from three set of calibration curves were listed in Table 3. Measurement accuracy (%) is calculated as the average of observed concentration divided by nominal concentration. In general, most B vitamers had accuracy within 20%, with a few exceptions such as nicotinic acid, 10-FDHF, 4-pyridoxic acid, cyanocobalamin, thiamine, pyridoxal and biotin at low concentrations (Q3). The first two vitamers, nicotinic acid and 10-FDHF, had Q3 below the limit of quantification (BLOQ) in all three set of calibration curves, while the last five had Q3 below the limit of quantification in set C calibration curves. Note that for calibration curves prepared in set C, since they went through the entire extraction procedure, some of the B vitamers had narrower linear range of quantification comparing to curves prepared in set A or B (Table 2), likely due to their instability under the extraction conditions. Examples include pyridoxal, cyanocobalamin, and 4-pyridoxic acid. Their signals had reduced significantly in set C comparing to that from set B, which led to low recovery of these B vitamers (Table 5). The instability of 10-FDHF was also reported previously (Zhang et al., 2018a). Their instability was also reflected from the measurement precision in set C (Table 4). Most B vitamers had intra- and inter-day precision within 15%, with exceptions from medium concentration of 10-FDHF in set C (21.3% and 29.2% for intra- and inter-day precision, respectively), and medium and high concentration of cyanocobalamin in set C (78.1% and 57.3% for intra-day, 75.0% and 33.0% for inter-day precision, respectively). The absolute recovery of each B vitamer was listed in Table 5. Recovery for most B vitamers ranged from 41.4% to 112.4%, with exceptions from the low concentration of six B vitamers (thiamine, nicotinic acid, 10-FDHF, 4-pyridoxic acid, biotin and pyridoxal) which were below the limit of quantification. The absolute recovery of cyanocobalamin in set C was very low (Table 5), with up to 95% of cyanocobalamin degraded (recovery ranged from 2.1% to 5.3%, with high variations). Similar observations were found for 4-pyridoxic acid, with 10.4% and 13.6% recoveries for the medium and high concentration QC samples, indicating its instability during the 4 h incubation at 37 °C. The instability of cyanocobalamin in the presence of ascorbate and other B vitamers was also reported previously (Ahmad, Qadeer, Zahid, Sheraz, Ismail, Hussain, & Ansari, 2014).

**Table 4:**
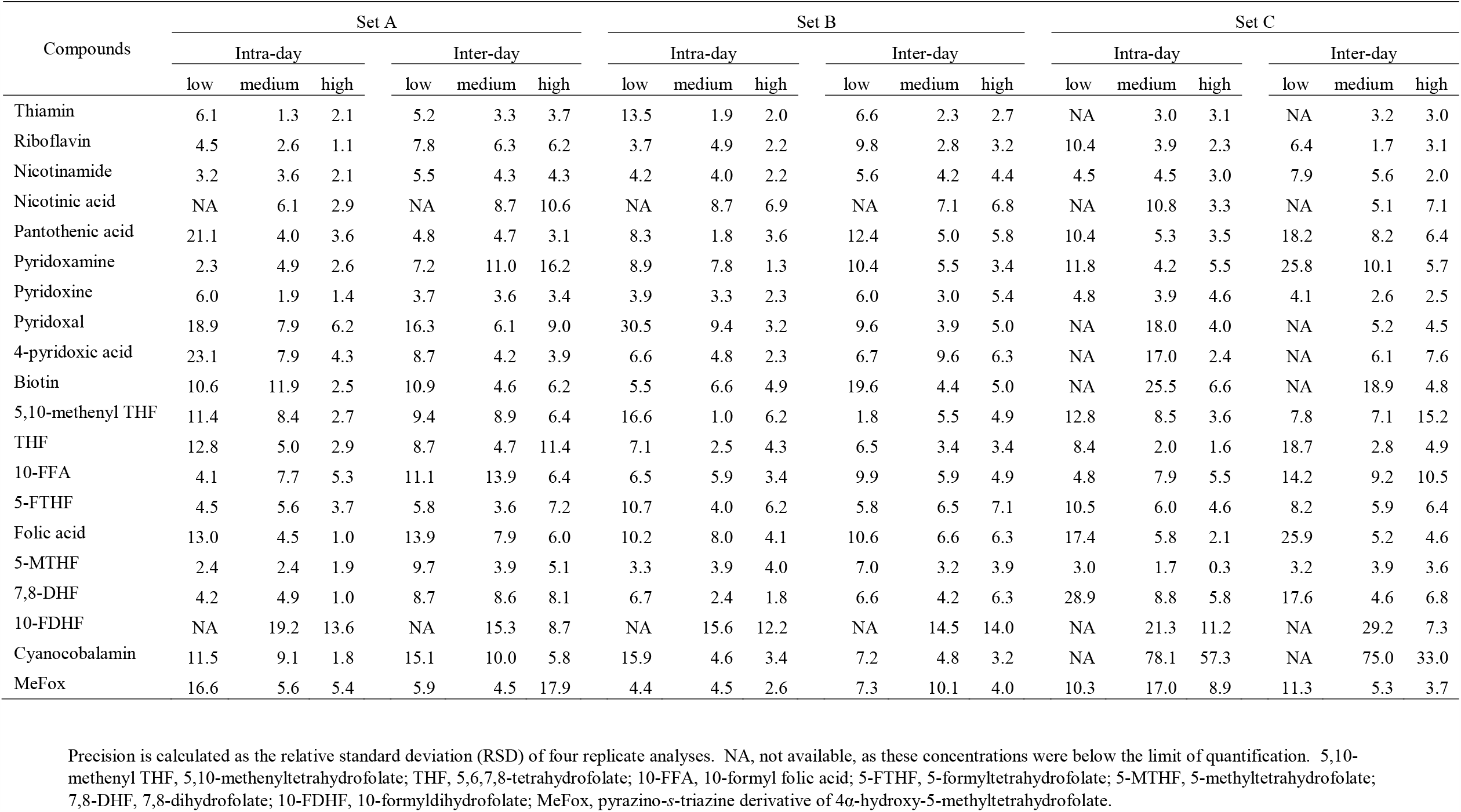
Intra-day (n=4) and inter-day (n=4) precision (%) of three QC standards from three sets of calibration curves.

**Table 5:** Absolute recovery (%) of three QC standards from three set of calibration curves. Data were from the average of six replicates (3 biological × 2 analytical replicates).

| Compounds | low | medium | high |
| --- | --- | --- | --- |
| Thiamin | NA | 103.4 ± 10.2 | 97.6 ± 4.4 |
| Riboflavin | 55.3 ± 21.5 | 57.5 ± 5.3 | 65.7 ± 7.0 |
| Nicotinamide | 54.1 ± 17.4 | 49.2 ± 15.8 | 52.0 ± 12.8 |
| Nicotinic acid | NA | 68.4 ± 11.6 | 68.0 ± 6.5 |
| Pantothenic acid | 61.8 ± 19.1 | 73.0 ± 14.5 | 82.2 ± 10.4 |
| Pyridoxamine | 61.5 ± 31.2 | 82.4 ± 6.5 | 76.4 ± 9.2 |
| Pyridoxine | 98.3 ± 2.1 | 99.2 ± 4.3 | 95.8 ± 5.0 |
| Pyridoxal | NA | 44.2 ± 28.2 | 47.1 ± 12.4 |
| 4-pyridoxic acid | NA | 13.6 ± 16.8 | 10.4 ± 20.0 |
| Biotin | NA | 104.2 ± 23.7 | 112.4 ± 19.3 |
| 5,10-methenyl THF | 76.7 ± 9.7 | 73.5 ± 14.1 | 78.5 ± 12.9 |
| THF | 59.8 ± 15.5 | 57.3 ± 13.2 | 56.4 ± 10.3 |
| 10-FFA | 61.1 ± 7.9 | 56.9 ± 8.6 | 55.0 ± 11.8 |
| 5-FTHF | 75.3 ± 13.3 | 79.5 ± 16.1 | 72.4 ± 13.9 |
| Folic acid | 41.4 ± 17.6 | 48.9 ± 7.5 | 45.8 ± 9.6 |
| 5-MTHF | 107.1 ± 9.8 | 97.1 ± 8.8 | 96.3 ± 9.8 |
| 7,8-DHF | 56.7 ± 17.8 | 59.0 ± 10.0 | 56.3 ± 15.1 |
| 10-FDHF | NA | 44.7 ± 23.6 | 63.3 ± 23.6 |
| Cyanocobalamin | NA | 2.1 ± 100.0 | 2.2 ± 85.0 |
| MeFox | 99.4 ± 7.3 | 95.1 ± 8.9 | 106.0 ± 3.3 |

Absolute recovery (%) is calculated from $C/B \times 100\%$ . C is denoted as the B vitamer peak area prepared in charcoal treated lentil matrix having gone through the entire extraction procedure (Set C), and B is denoted as the B vitamer peak area prepared in treated matrix and directly injected for LC-MS (Set B). NA, not available, as these concentrations were below the limit of quantification. 5,10-methenyl THF, 5,10-methenyltetrahydrofolate; THF, 5,6,7,8-tetrahydrofolate; 10-FFA, 10-formyl folic acid; 5-FTHF, 5-formyltetrahydrofolate; 5-MTHF, 5-methyltetrahydrofolate; 7,8-DHF, 7,8-dihydrofolate; 10-FDHF, 10-formyldihydrofolate; MeFox, pyrazino-s-triazine derivative of 4α-hydroxy-5-methyltetrahydrofolate.

Matrix effect was also investigated for the original matrix solution (1×), and five times diluted matrix (5×), as shown in Table 6. With the normalization of isotope labeled internal standards (ISs), most B vitamers had matrix effects between 91.1% to 108.8% in original matrix (1× matrix), with the exception of pyridoxamine (72.9%), 4-pyridoxic acid (77.7%) and 7,8-DHF (74.8%), which had lower matrix effects, and 10-FDHF (126.1%), which had higher matrix effect. Dilution of treated matrix 5 times with extraction buffer (50 mM MES, pH 6.0), the matrix effects of most B vitamers was within acceptable range (85.4% to 108.4%)without internal standard normalization, except for pantothenic acid (72.2%), 5,10-methenyl THF (81.2%), AND 7,8-DHF (83.9%). With internal standard normalization, the matrix effects of all B vitamers fell into acceptable range (87.3% to 103.2%). Similar observation has been reported for folate quantification with 5× diluted matrix (Zhang et al., 2019). To represent the real sampling conditions, similar concentration of treated matrix blank (as samples) should be used for preparing calibration curves.

**Table 6:** Matrix effect (%) of each B vitamer from three sets of calibration curves. Data were from the average of six replicates (3 biological × 2 analytical replicates).

| Compounds | 1× matrix | | 5× matrix | |
| --- | --- | --- | --- | --- |
|  | without IS | with IS | without IS | with IS |
| Thiamine | 134.3 | 100.6 | 108.4 | 99.2 |
| Riboflavin | 91.8 | 98.8 | 93.3 | 96.7 |
| Nicotinamide | 78.2 | 96.8 | 85.4 | 95.0 |
| Nicotinic acid | 90.3 | 108.8 | 85.7 | 95.5 |
| Pantothenic acid | 105.4 | 90.7 | 72.2 | 101.1 |
| Pyridoxamine | 75.3 | 72.9 | 89.3 | 87.3 |
| Pyridoxine | 100.3 | 98.4 | 99.8 | 97.6 |
| Pyridoxal | 94.6 | 91.1 | 93.8 | 91.6 |
| 4-pyridoxic acid | 79.7 | 77.7 | 91.9 | 89.4 |
| Biotin | 79.6 | 97.6 | 86.0 | 96.5 |
| 5,10-methenyl THF | 60.6 | 96.9 | 81.2 | 89.4 |
| THF | 97.2 | 97.7 | 87.2 | 96.2 |
| 10-FFA | 87.4 | 101.8 | 89.5 | 98.8 |
| 5-FTHF | 84.6 | 98.4 | 85.7 | 96.2 |
| Folic acid | 93.0 | 93.2 | 89.8 | 94.8 |
| 5-MTHF | 62.1 | 98.4 | 88.6 | 98.0 |
| 7,8-DHF | 75.4 | 74.8 | 83.9 | 89.3 |
| 10-FDHF | 123.3 | 126.1 | 92.2 | 103.2 |
| Cyanocobalamin | 92.0 | 99.4 | 92.3 | 95.7 |
| MeFox | 118.7 | 101.9 | 88.8 | 99.5 |

IS, internal standard; Matrix effect (%) without IS is calculated as $B/A \times 100\%$ . A is the B vitamer peak area prepared in MES buffer (from set A), and B is the B vitamer peak area prepared in blank lentil matrix (from set B). Results with IS are calculated as $[(B/IS_B)/(A/IS_A)] \times 100\%$ , in which $IS_B$ and $IS_A$ are peak areas of isotope labeled standards from set B and set A, respectively. 5,10-methenyl THF, 5,10-methenyltetrahydrofolate; THF, 5,6,7,8-tetrahydrofolate; 10-FFA, 10-formyl folic acid; 5-FTHF, 5-formyltetrahydrofolate; 5-MTHF, 5-methyltetrahydrofolate; 7,8-DHF, 7,8-dihydrofolate; 10-FDHF, 10-formyldihydrofolate; MeFox, pyrazino-*s*-triazine derivative of 4α-hydroxy-5-methyltetrahydrofolate.

### 3.4. B vitamin concentration change during lentil seed germination

Germinated seeds have become attractive functional foods in recent years (Benincasa, Falcinelli, Lutts, Stagnari, & Galieni, 2019; Idowu, Olatunde, Adekoya, & Idowu, 2019), due to their increased nutritional value compared to dry seeds, such as the increased levels of phytochemicals and bioactive peptides, accompanied by a reduction in antinutritional compounds such as phytate, tannin and trypsin inhibitor. In the current work, a zero-tannin lentil experimental line was selected for B vitamin quantification during seed germination. For nicotinic acid, nicotinamide and pantothenic acid, due to their high concentration in germinated seeds, a 5× dilution of the original extracts was needed for accurate measurement. Therefore, calibration curves prepared in 5× diluted treated matrix were used for their quantification. For the rest of B vitamers, the original extracts (1×) was used for quantification, together with calibration curves prepared in 1× treated matrix. The individual B vitamer concentrations at each day of germination were listed in supplementary Table S1, and the concentration changes of each B vitamin group and the total B vitamins were shown in Figure 2. In dry lentil seeds (day 0), B5 (pantothenic acid, 13.4 ± 0.85 µg/g) and B3 (12.5 ± 1.17 µg/g) had the highest concentration, followed by B1 (5.33 ± 0.05 µg/g) and B6 (3.4 ± 0.12 µg/g). B8 (biotin) and B12 (cyanocobalamin) were not detected either in dry seeds, or during germination. In addition, three other B vitamers, 4-pyridoxic acid (B6), 10-FDHF (B9) and 7,8-DHF (B9), were not detected in these samples, therefore they were not listed in Table S1.

**Figure 2.**
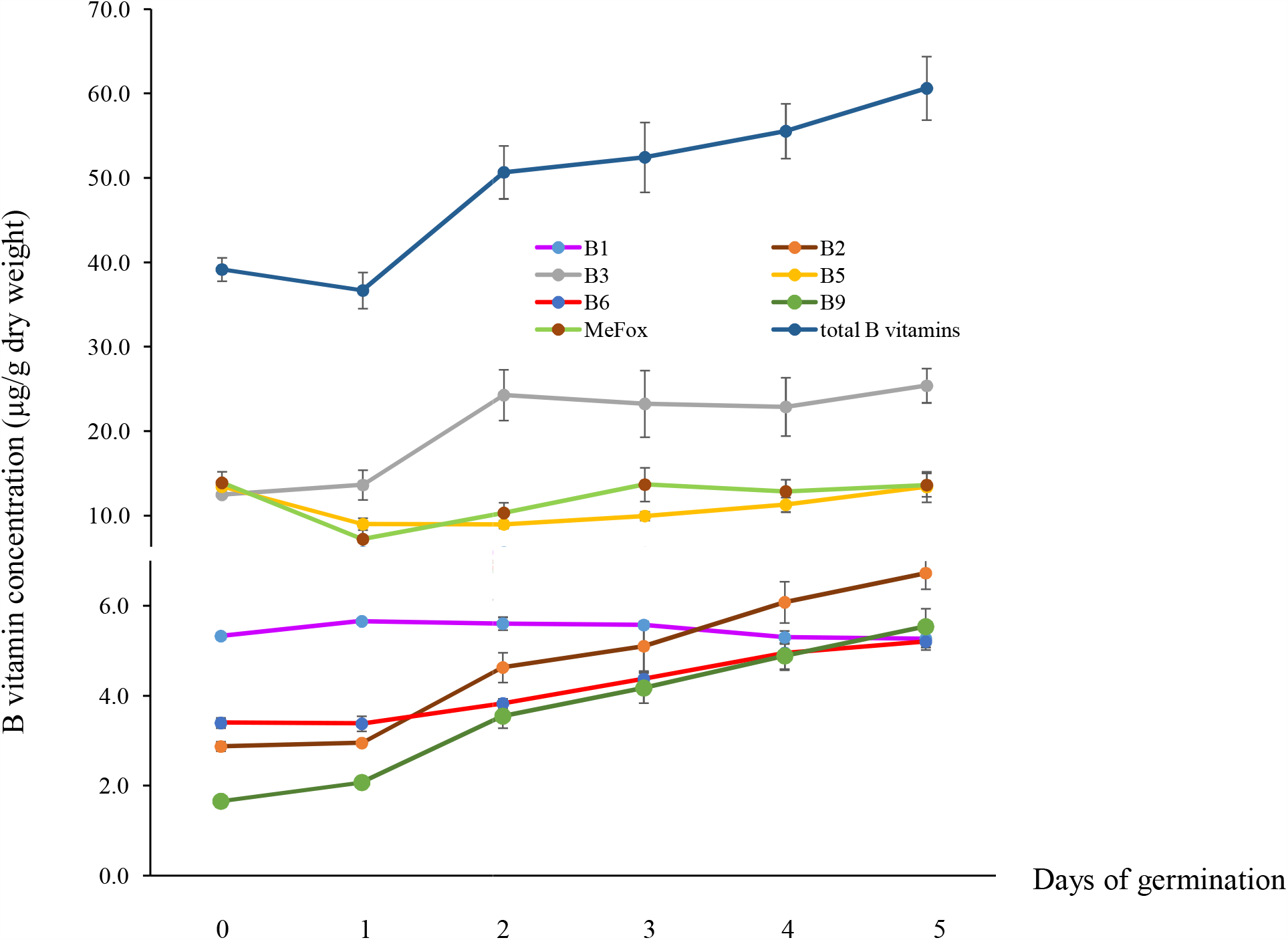
B vitamin concentration change during seed germination (from a Zt-lentil experimental line). B1 (purple), B2 (red), B3 (grey), B5 (orange), B6 (blue), B9 (green), and total B vitamin (dark red). Error bars were the standard deviation from six replicates (3 biological × 2 analytical replicates).

During seed germination, total B vitamin concentration declined on day 1 (from 39.2 to 36.7 µg/g), and gradually increased from day 2 to day 5 (from 50.7 to 60.6 µg/g). By day 2, the total B vitamins in lentil sprouts were significantly higher than the dry lentil seeds (P<0.01), and by day 5, the total B vitamin increased 1.5-fold compared to the dry seeds. For individual B vitamin groups, B5 (pantothenic acid) showed a dramatic decrease (−33.1%) during early germination (days 1 and 2), and started to increase on day 3. By day 5 its concentration was close to the original concentration in dry seeds. B1 (thiamin) concentration was almost consistent during germination, with 5.33 µg/g in dry seeds, and 5.27 µg/g on day 5 of germination. Compared with dry seeds, B3 and B6 started to increase on day 2, and reached their maximum on day 5 (up to 104% and 53.7% increase on day 5, respectively). B9 concentration started to increase on day 1 (1.25 fold) of germination, and reached to maximum on day 5 (3.34 fold). MeFox, being an oxidation product of 5-MTHF (Ringling & Rychlik, 2017), showed a large decrease on day 1 (−48.1%), and then gradually increased from day 2. By day 5 its concentration was almost the same as in dry seeds (13.9 µg/g in dry seeds, and 13.6 µg/g in germinated sprouts). Riboflavin started to increase on day 2 (1.6× fold), and on day 5, its concentration increased up to 2.3× fold comparing to dry seeds.

The fact that thiamin had no significant change during germination was also reported in lentil (Kylan & McCready, 1975; Urbano et al., 1995; Vidal-Valverde et al., 2002), pea and beans (Urbano et al., 2005; Vidal-Valverde et al., 2002), corn and oats (Cheldelin & Lane, 1943). Riboflavin was also reported to increase during seed germination from pulses, cereal and oilseeds (Burkholder & McVeigh, 1942; Cheldelin & Lane, 1943; Kavas & El, 1991; Urbano et al., 1995; Vidal-Valverde et al., 2002). The increase of niacin (B3) in legume (Prodanov, Sierra, & Vidal-Valverde, 1997), nicotinic acid (B3) and pyridoxin (B6) in lima bean and cotton (Cheldelin & Lane, 1943) after germination were reported, and folate concentration increase during germination were reported in rye (Kariluoto et al., 2006), soybean and mung bean (Shohag, Wei, & Yang, 2012). This suggested that germination is an economic and effective way to improve the nutrition value of edible seeds, including lentil (Idowu et al., 2019).

## 4. Conclusions

A validated B vitamin quantification protocol was reported, covering 20 B vitamers from eight B vitamin groups. A 5-min UPLC-SRM MS method was optimized which provided good peak separation and short analysis time, thus making it suitable for high throughput analysis. Although 4-pyridoxic acid (B6) and cyanocobalamin (B12) (Ahmad et al., 2014) were not stable under current extraction condition after 4 h incubation, this UPLC-SRM MS method is still applicable to their quantifications from B vitamin fortified foods or energy drinks (Bendryshev et al., 2010), in which quick extraction is possible and sample degradation could be minimized. To our knowledge, this is the first systematic study which can simultaneously quantify 20 B vitamers with a short separation time. The calibration curve and sensitivity data of each B vitamer was determined, and this method was applied to B vitamin quantification from germinated lentil seeds. The use of three enzymes (acid phosphatase, β-glucosidase, and rat serum) in a single extraction step with 4 h incubation was determined to be sufficient to convert B vitamins into their free forms. Within comparison to other B vitamin extraction procedures which required up to 18 h incubation (Ndaw et al., 2000; Nurit et al., 2015), the current approach enables sample extraction from ground seeds to data collection to be conducted within one work day. After seed germination, lentil sprouts could provide higher concentration of B vitamins than dry seeds, which makes it an attractive source of functional food.

## Author contributions

Haixia Zhang: supervision, conception and design of study, conducted experiment, data acquisition and analysis, drafting the manuscript; Devini De Silva and Dilanganie Dissanayaka, conducted experiment; Thomas Warkentin, funding acquisition; Albert Vandenberg: conception and design of study, funding acquisition. All authors reviewed the manuscript critically for important intellectual content.

## Acknowledgements

The authors acknowledge the technical assistance from Jeremy Marshall, and the Pulse Crop Field Laboratory technical staff at the University of Saskatchewan for their assistance with plant and seed production.

## Supporting information description

### Supplementary data

Supplementary material related to this article can be found, in the online version, at doi:

---

**Table S1:** 17 B vitamers and MeFox concentrations in an experimental lentil line, during seed germination (µg/g of dry weight, average ± SD). Data were from the average of six replicates (3 biological × 2 analytical replicates).

| Compounds | Day 0 | Day 1 | Day 2 | Day 3 | Day 4 | Day 5 |
| --- | --- | --- | --- | --- | --- | --- |
| Thiamine | 5.33 ± 0.05 | 5.66 ± 0.04 | 5.61 ± 0.14 | 5.58 ± 0.08 | 5.30 ± 0.14 | 5.27 ± 0.25 |
| Riboflavin | 2.87 ± 0.11 | 2.95 ± 0.06 | 4.63 ± 0.33 | 5.10 ± 0.55 | 6.08 ± 0.46 | 6.73 ± 0.36 |
| Nicotinamide | 8.48 ± 0.86 | 6.14 ± 1.83 | 6.66 ± 0.75 | 2.34 ± 1.85 | 0.65 ± 0.24 | 1.01 ± 0.30 |
| nicotinic acid | 4.00 ± 0.85 | 7.01 ± 1.37 | 19.2 ± 1.42 | 20.6 ± 1.73 | 23.5 ± 2.04 | 24.3 ± 2.00 |
| pantothenic acid | 13.4 ± 0.85 | 8.99 ± 0.70 | 8.97 ± 0.46 | 9.95 ± 0.52 | 11.3 ± 0.87 | 12.7 ± 0.92 |
| Pyridoxamine | 0.23 ± 0.01 | 0.09 ± 0.01 | 0.09 ± 0.00 | 0.09 ± 0.01 | 0.09 ± 0.01 | 0.09 ± 0.01 |
| Pyridoxine | 2.57 ± 0.07 | 2.39 ± 0.13 | 2.77 ± 0.12 | 3.33 ± 0.15 | 4.01 ± 0.34 | 3.89 ± 0.32 |
| Pyridoxal | 0.59 ± 0.05 | 0.90 ± 0.13 | 0.97 ± 0.09 | 0.95 ± 0.13 | 0.86 ± 0.06 | 1.23 ± 0.23 |
| 5,10-methenyl THF | 0.06 ± 0.01 | 0.04 ± 0.01 | 0.12 ± 0.03 | 0.13 ± 0.06 | 0.14 ± 0.06 | 0.10 ± 0.03 |
| THF | 0.26 ± 0.04 | 0.09 ± 0.03 | 0.14 ± 0.03 | 0.15 ± 0.03 | 0.28 ± 0.07 | 0.23 ± 0.04 |
| 10-FFA | 0.09 ± 0.01 | 0.04 ± 0.00 | 0.03 ± 0.00 | 0.03 ± 0.01 | 0.05 ± 0.01 | 0.10 ± 0.05 |
| 5-FTHF | 0.41 ± 0.01 | 0.08 ± 0.02 | 0.06 ± 0.01 | 0.09 ± 0.04 | 0.08 ± 0.01 | 0.09 ± 0.01 |
| Folic acid | 0.01 ± 0.00 | 0.01 ± 0.01 | 0.02 ± 0.01 | 0.03 ± 0.01 | 0.07 ± 0.03 | 0.28 ± 0.25 |
| 5-MTHF | 0.83 ± 0.03 | 1.82 ± 0.06 | 3.18 ± 0.25 | 3.74 ± 0.31 | 4.28 ± 0.20 | 4.74 ± 0.18 |
| Total B vitamins | 39.2 ± 1.39 | 36.7 ± 2.14 | 50.7 ± 3.14 | 52.5 ± 4.13 | 55.6 ± 3.24 | 60.6 ± 3.77 |
| MeFox | 13.9 ± 1.30 | 7.21 ± 1.38 | 10.3 ± 1.23 | 13.7 ± 2.00 | 12.9 ± 1.38 | 13.6 ± 1.38 |

SD, standard deviation; 5,10-methenyl THF, 5,10-methenyltetrahydrofolate; THF, 5,6,7,8-tetrahydrofolate; 10-FFA, 10-formyl folic acid; 5-FTHF, 5-formyltetrahydrofolate; 5-MTHF, 5-methyltetrahydrofolate; MeFox, pyrazino-*s*-triazine derivative of 4α-hydroxy-5-methyltetrahydrofolate.

